# Recurring Transient Brain-Wide Co-Activation Patterns from EEG Spatially Resembling Time-Averaged Resting-State Networks

**DOI:** 10.1101/2025.06.26.661612

**Authors:** KC K. Nkurumeh, Han Yuan, Lei Ding

**Affiliations:** Stephenson School of Biomedical Engineering, University of Oklahoma, Norman, USA; Institute for Biomedical Engineering, Science, and Technology, University of Oklahoma, Norman, USA

**Keywords:** Co-activation pattern, electroencephalography, resting-state network, independent component analysis, temporal dynamics, functional connectivity

## Abstract

It has long been established that human brains remain functionally active at rest, as demonstrated with the discovery of resting-state networks (RSNs) underlying spontaneous neural activity. Recent studies suggest that classical RSNs estimated from functional magnetic resonance imaging (fMRI) data using time-domain functional connectivity measures might be driven by recurring point-process events. Due to the slow hemodynamic response, fMRI is not able to reveal such point-processes at the timescale of neuronal events while electroencephalography (EEG) holds the promise due to its millisecond temporal resolution and successful reconstruction of fMRI-like RSNs from EEG. The present study reported a set of recurring transient (<100 milliseconds) cortical co-activation patterns (CAPs) derived from resting-state EEG using a clustering algorithm with spatial-domain measures (i.e., k-means). Our results indicate that this set of CAPs exhibit strong spatial correspondence with known RSNs, not only those derived from the same EEG data using time-domain measures (i.e., independence), but also those from fMRI literature, covering visual, auditory, motor, limbic, high-order, and default mode networks. CAPs exhibit the properties of hemispheric symmetry, spatially separatable sub-systems, and intersubject variability gradient across functional systems, which have all been observed in classical RSNs. In terms of differences between CAPs and RSNs beyond their timescales, CAPs demonstrate greater intersubject reproducibility of spatial patterns compared to their time-averaged RSNs counterparts. These findings suggest that classical RSNs might be driven by recurring transient neuronal activations captured in CAPs. More importantly, CAPs can reveal much fast dynamics of such brain-wide networked neuronal activations (e.g., different CAPs exhibit significantly different occurrences and lifetimes) and benefit from their great intersubject reproducibility, thus underscoring their potential to advance our understanding on neuronal mechanisms of spontaneous large-scale brain activation phenomena.

## 1. Introduction

Human brains are organized into multi-layer networks, and such hierarchical structures serve as neural substrates for distinct brain functions (Biswal et al., 1995; Fox & Raichle, 2007), especially for high-level functions such as cognition (Bressler & Menon, 2010). These networks can be modulated by tasks (Betti et al., 2013; Krienen et al., 2014) while their intrinsic natures are present even without explicit tasks (Cole et al., 2014). Such intrinsic phenomena have been widely captured in neuroimaging data, e.g., functional magnetic resonance imaging (fMRI), electroencephalography (EEG), magnetoencephalography (MEG), and functional near-infrared spectroscopy (fNIRS). Their spatial constructions are obtained via calculating functional connectivity (FC) measures among signals recorded between different locations, known as resting-state networks (RSNs) (Jafri et al., 2008). The properties of these RSNs have shown significant behavioral correlates (Rosenberg et al., 2016) and have been associated with clinical abnormalities in major neuropsychiatric disorders (Fox & Greicius, 2010; Friston, 2011).

While RSNs were firstly reconstructed statically from entire recordings of a few minutes of fMRI data (therefore time-averaged) using FC measures including pairwise statistical metrics, e.g., correlation (Biswal et al., 1995; Fox et al., 2005) or data-driven methods, e.g., independent component analysis (ICA) (Beckmann et al., 2005; Fox & Raichle, 2007), later studies have demonstrated that RSNs show dynamics over the time scale of minutes in fMRI data. Some recent studies further indicate that fMRI-based network dynamics at much shorter temporal scales (i.e., ∼22.5 seconds) can be used in tracking cognition while performing multiple tasks (Gonzalez-Castillo et al., 2015), which suggest potential sub-minute FC dynamics in RSNs. These time-domain FC measures, e.g., correlation (De Pasquale et al., 2010) and independence (Li et al., 2018; Yuan et al., 2012), and their corresponding sliding-time versions (Shou et al., 2020), have been utilized on EEG/MEG signals and successfully revealed both static and dynamic RSNs. Due to their millisecond (ms) temporal resolutions, EEG/MEG signals hold the potential in revealing RSN dynamics at the timescale of neuronal events (i.e., tens to hundreds of milliseconds). The adoptions of novel measures, e.g., spatial-domain measures (as opposed to time-domain FC measures) (X. Liu & Duyn, 2013), advanced methods, e.g., Hidden Markov Methods (HMM) (Baker et al., 2014; Vidaurre et al., 2017), and other ad-hoc algorithms (Gu et al., 2021; Matsui et al., 2016) have led to the identifications of several new large-scale patterns in both fMRI and EEG/MEG at the temporal resolution of individual timeframes (only limited by sampling rates of corresponding neuroimaging modalities). While most of these large-scale patterns, e.g., propagational waves (Matsui et al., 2016), co-activation patterns (CAPs) (X. Liu & Duyn, 2013), and HMM brain states (Baker et al., 2014), are typically not named as RSNs as they are not constructed using time-domain FC measures, they share the similar characteristic brain-wide spatial distributions as RSNs but usually are of transient and recurring nature. In particular, such EEG/MEG-based large-scale patterns exhibit their lifetimes at the time scale of neuronal events, i.e., tens to hundreds of milliseconds (Ding et al., 2022; Vidaurre et al., 2018).

Such a collection of large-scale patterns spanning the timescales from ten/hundred milliseconds to several minutes and observable in various contrast signals (hemodynamic and electrical) provide the unique opportunities in understanding neuronal mechanisms of classical time-averaged RSNs obtained using time-domain FC measures (Preti et al., 2017). Firstly, the spatial similarity between EEG- and fMRI-derived RSNs suggests a possible shared neuronal basis among them (Mantini et al., 2007; Yuan et al., 2012; Yuan et al., 2016). Secondly, recurring fMRI CAPs spatially resemble fMRI RSNs, which may contribute to the emergence of RSNs observed over longer time windows (X. Liu et al., 2013). Thirdly, the detection of both CAPs and RSNs in human and animal models points to their evolutionary relevance as functional motifs (Belloy et al., 2018). Lastly, while both EEG RSNs and EEG CAPs have been reported and some spatial similarities have been discussed (Ding et al., 2022), current evidence suggests more discrepancies (Shou et al., 2022) in their spatial correspondence than those seen between fMRI RSNs and fMRI CAPs. It is therefore of interest to study whether there are another set of EEG CAPs that exhibit more similarities to EEG RSNs, which can fill the missing link between large-scale transient patterns and large-scale time-averaged patterns in electrical signals as the similar link has been observed in hemodynamic signals.

In the present study, we reported a set of EEG-derived CAPs that resemble time-averaged RSNs identified using traditional ICA approaches. These CAPs were extracted using the k-means clustering with correlation as the distance measure from cortical envelope data that were inversely reconstructed from surface EEG. The outcome from the present study reveals several important phenomena. Firstly, RSN-like patterns (i.e., CAPs) could be identified at the timescale of typical neuronal events (<100 ms), which is several orders shorter than time windows typically used for calculating time-averaged RSNs. Secondly, these CAPs occur repeatedly within participants and are reproducible across participants, often with greater consistency than their ICA-derived counterparts. Thirdly, several canonical distinctive features observed in RSNs are also observed in CAPs, including hemisphere symmetry, intersubject variability gradient from low-level perceptual systems to high-level functional systems, and existence of both default mode network (DMN) and task positive networks (TPNs). Finally, CAP spatial patterns remain stable across a range of values for a key method parameter, demonstrating their robustness to methodological factors. These observations support the hypotheses that there are large-scale transient patterns matched to large-scale time-averaged patterns in resting human brains.

## 2. Materials and Methods

### 2.1 Data Acquisition and Preprocessing

The study was approved by IRB at the University of Oklahoma Health Sciences Center (OUHSC) and written informed consent was obtained from all participants before data acquisition. EEG and MRI data from 34 healthy participants (age = 24 ± 5 years, 9 females, absence of known brain diseases or neurological complications) were acquired. Participants were instructed to remain seated upright with eyes closed during EEG recordings. Resting-state EEG data were recorded for 10 minutes, at the sampling rate of 1000 Hz, using a 128-channel Amps 300 amplifier (Electrical Geodesics Inc., OR, USA) in each participant. Positions of EEG sensors along with three landmarks (nasion, left and right pre-auricular points) were digitized by the Polhemus Patriot system. Structural MRI (sMRI) covering full head was collected for each participant on a GE MR750 scanner at OUHSC MRI facility, using the GE BRAVO sequence: FOV = 240 mm, axial slices per slab = 180, slice thickness = 1 mm, image matrix = 256 × 256, TR = 8.45 ms, TE = 3.24 ms.

All EEG preprocessing was performed using the EEGLAB toolbox (Delorme & Makeig, 2004) separately for individuals. A notch filter of 58-62 Hz and a bandpass filter of 0.5-100 Hz were applied. Noisy channels were automatically identified using the FASTER plugin (Nolan et al., 2010) for EEGLAB and interpolated using data from their neighboring channels. ICA was then applied to identify and remove artifactual components related to cardiac, ocular, or muscular activity by visual inspection of ICs’ topographies, time courses, and spectral properties (Chaumon et al., 2015). Finally, EEG data were downsampled to 250 Hz and referenced to the common average. Note that no time segments of EEG data were removed in order to preserve continuity of data.

### 2.2 Reconstructing Cortical Source Tomography

Cortical source imaging (Li et al., 2018) was used to construct current source distributions over the cortical surface from scalp EEG. Firstly, a boundary element model was generated for each participant. Freesurfer (Fischl, 2012) was used to segment and extract the surface boundaries of the scalp, skull, and brain and the surface boundary between white and grey matters from participant’s sMRI data to create individualized volume conduction models and cortical current density (CCD) source models, respectively. Next, each surface of the volume conduction model was tessellated into 10,242 nodes and 20,480 triangular elements, while the CCD surface was tessellated into 20,484 nodes and 40,960 triangular elements. The nodes along the medial wall comprising the corpus callosum, basal forebrain, and hippocampus were excluded, therefore reducing the total number of source nodes to 18,715. Each node of the CCD surface was assigned a dipole vector to represent local neuronal currents with its direction defined as the normalized sum of the normal vectors of all triangles sharing the node. The electrical conductivities of the scalp, skull, and brain of the volume conduction model were assigned as 0.33/Ωm, 0.0132/Ωm, and 0.33/Ωm, respectively (Lai et al., 2005). The locations of EEG sensors were registered to the scalp surface by aligning three landmarks from both EEG and sMRI recordings.

Based on the volume conduction and CCD models, the boundary element method (Hamalainen & Sarvas, 1989) was used to build the forward relationship: ***Φ***(***t***) = ***L*** · ***S***(***t***), in which ***L*** is an 128×18715 lead field matrix representing the contribution of each cortical unity dipole on EEG signals recorded at all sensors and ***Φ***(***t***) and ***S***(***t***) are the EEG recordings and dipole source amplitude as functions of time. Based on the forward relationship, the minimum-norm estimate was used to calculate cortical source amplitudes: ***S***(***t***) = ***L***^***T***^ · (***L*** · ***L***^***T***^ + ***λ***(***t***) · ***I***)***^−1^*** · ***Φ***(***t***) (Hämäläinen & Ilmoniemi, 1994), in which *I* is the identity matrix, and ***λ*** is the regularization parameter for each time instance selected via the generalized cross validation method (Golub et al., 1979). To control the quality of cortical source estimates, values of ***λ*** outside of three standard deviations of ***λ*** values from all time instances were considered as outliers and interpolated with neighboring values. Based on the adjusted ***λ*** values, cortical current source tomography was constructed for individual participants as functions of time.

### 2.3 Estimating Time-averaged RSNs with Group-level ICA and Participant-level Statistical Regressions

To estimate time-averaged RSNs, we performed the group-level time-frequency ICA on the alpha band (8– 12 Hz) of preprocessed EEG data and then statistical regression analysis to obtain individual representations of RSNs with both individualized spatial patterns and associated time courses (Ding et al., 2022). Alpha band data was chosen because it is the dominant rhythmic neural oscillation in resting human brains (Kirschfeld, 2005). To perform the group-level ICA, each participant’s data was converted to z-scores channel-wise to remove inter-participant variabilities. Short-time Fourier transform was used to calculate the complex spectral time series representation of time-domain individual EEG data (0–100 Hz, 1 Hz steps, 1 second window, no overlap). Complex spectral EEG data in the range of 8–12 Hz was then temporally concatenated across all individuals subject to the time-frequency ICA (Bingham & Hyvärinen, 2000; Shou et al., 2012) using various numbers of ICs (i.e., 25, 36, 48, and 64) where the maximal number investigated (i.e., 64) was recommended as more ICs (>64) tended to generate focal and regional subnetworks (Smith et al., 2009). Results from 48 ICs were selected for the subsequent analysis to minimize model complexity while retaining sufficient number of valuable components. To obtain IC time courses, original alpha-band EEG data were projected using the demixing matrix obtained in the group-level ICA. ICs showing neural activation characteristics in both their spatial and spectral patterns comparable to neuronal ICs reported in literature (Brookes et al., 2011; Shou et al., 2012, 2020; Yuan et al., 2016) were selected as the scalp-level representations of time-averaged RSNs, which led to the selection of 29 neuronal ICs out of total 48 ICs (see Supplementary Fig. 1).

Cortical representations of time-averaged RSNs were estimated via the statistical regression analysis (Ding et al., 2022) between the time courses of the selected neuronal ICs and the activation time courses of cortical dipole sources (Section 2.2). Firstly, cortical dipole time courses were bandpass filtered to the alpha band (i.e., 8–12 Hz). Instantaneous amplitudes of the time courses of both neuronal ICs and dipoles were then estimated using the Hilbert transform (Baker et al., 2014; Coquelet et al., 2022), and converted to z-scores timewise. Thirdly, a regression was performed for each cortical dipole with the dipole amplitude time course as the response data and the amplitude time courses of all 48 ICs (29 neuronal and 19 non-neuronal ICs) as the regressors. Regression in this manner resulted in participant-level spatial maps of ß-values for each regressor (i.e., each IC). The group-level cortical RSNs were calculated as the average of corresponding participant-level cortical tomography from individual ICs.

### 2.4 Estimation of CAPs: Clustering

To estimate CAPs, cortical dipole time courses were subject to k-means clustering at the resolution of individual timeframes. Instantaneous amplitudes of cortical dipole time courses from individual participants that were bandpass filtered to the alpha band (Section 2.3) were used. A dimension reduction from 18,715 cortical sources to 200 regions of interest (ROIs) was performed using a well-established atlas for human cortex (Schaefer et al., 2018). The ROI-based cortical source amplitude data were calculated as the average source amplitude of all nodes within an ROI at any given time point. The ROI amplitude data were converted to z-scores timewise to minimize inter-participant variabilities that allowed all ROIs to be equally weighted in the k-means algorithm. Then, the ROI-based z-score time courses from all participants was concatenated as the input to the k-means algorithm (Lloyd, 1982) using correlation as the distance measure for the number of clusters (k) ranging from 10 to 20. The outcome of the algorithm labels each timeframe to a cluster out of total k clusters based on similarities among individual timeframe data. The participant-level spatial patterns of each CAP (i.e., a cluster) was constructed via averaging of all timeframe data labeled for the cluster in individual participants. The group-level CAPs were calculated by averaging all participant-level spatial maps of the corresponding CAPs. Normalized cortical source data were used to obtain these spatial maps as opposed to the ROI data to maintain high spatial resolutions. Temporal properties of CAPs were defined in two measures, i.e., lifetime and occurrence rate. An occurrence of a CAP was defined as multiple consecutive timeframes that were labeled toward the same CAP, and the occurrence rate of a CAP was its total number of occurrences divided by the entire time of recording from each participant. Lifetime of a CAP was calculated for each occurrence as the total time of consecutive timeframes of one occurrence. These two temporal measures were calculated in individual participants and then averaged for group-level statistics. In the subsequent analysis, results of 20 CAPs (k = 20) were used.

### 2.5 Matching and Comparisons of CAPs and Time-Averaged RSNs

Obtained CAPs were named according to the resemblance of their spatial patterns to known anatomical regions and time-averaged RSNs reported in literature (Beckmann et al., 2005; Thomas Yeo et al., 2011). Each CAP was either resolved as a hemisphere-bilateral pattern or as one of a hemisphere-symmetric CAP pair. These symmetrically paired CAPs were named with “left (L)” or “right (R)” after their anatomical names to indicate their dominant side. To facilitate comparisons between large-scale transient patterns, i.e., CAPs, and time-averaged RSNs, i.e., ICs, the 20 CAPs were matched one-to-one with the 29 neuronal ICs through visual inspections. However, several CAPs showed bilateral patterns while their corresponding ICs were unilateral. In these cases, a single CAP was matched with the pair of hemisphere-symmetric ICs. The same one-to-two matching was applied if an IC was bilateral and its corresponding CAPs were unilateral.

To validate the matched outcomes, Pearson spatial correlation was computed between all matched pairs of a CAP and an IC. If there were multiple ICs that expressed similar spatial patterns to one CAP, only the IC with the highest regression beta value were selected, which reduced the number of matched ICs to 17. For a bilateral CAP matched with two hemisphere-symmetric ICs (or vice versa), the single-hemisphere spatial correlations were calculated on left hemisphere and right hemisphere separately with respect to the corresponding hemisphere-dominant IC and then averaged over two calculations. All spatial correlation values were then converted to z-scores using the Fisher transform to normalize their distributions to build the final confusion matrix of 20 CAPs and 17 RSNs. One-way ANOVA was performed to evaluate the difference in spatial correlation values from matched and unmatched CAP-RSN pairs.

Beyond comparing spatial patterns of CAPs directly to RSNs, the hemispheric symmetries for all CAPs were also evaluated. To assess the hemispheric symmetry of CAPs and RSNs, spatial patterns were averaged into 148 ROIs (74 matching ROIs in each hemisphere) using a well-established cortical surface atlas (Destrieux et al., 2010). The hemispheric symmetry index was calculated as the Pearson correlation between the original ROI-based spatial pattern and their mirrored ones (ROIs values swapped between the left and right hemispheres). For the unilateral CAP pairs, the original and mirrored ones referred to one of the pairs. For the bilateral CAPs, the mirrored one was from itself.

## 3. Results

### 3.1 Transient CAPs spatially resembling time-averaged RSNs

According to the naming protocol in Section 2.5, 20 CAPs are categorized into three groups: Sensorimotor (SM) CAPs, each mainly covering cortical areas responsible for sensory or motor functions; High-Order (HO) CAPs, each mainly covering either frontal, temporal, and/or parietal cortical areas; and Other CAPs, each covering either distributed cortical areas or limbic areas. Out of the total 20 CAPs, 15 CAPs are successfully matched with RSNs (Fig. 1A). From the perspective of RSNs, the majority of ICs were also matched with CAPs (19 out of total 29 neuronal ICs). It is noted that the remaining 10 unmatched neuronal ICs in general exhibit lower regression beta values (indicating lower contributions to data variances) than matched ones (Supplementary Fig. 2B), which are of statistical significance (*p<0.001*, *corrected*, Supplementary Fig. 2D). Fig. 1A illustrates the confusion matrix of spatial correlation values between 20 CAPs and 17 RSNs (after removing duplicated matches). Among the combinations of 20 CAPs and 19 RSNs, the matched pairs (elements emphasized with black boxes) in general have higher z-scored correlation values than the non-matched pairs (unboxed elements). Statistically, spatial correlation values from all matched pairs (the blue bar in Fig. 1B) were found to be significantly higher (*p<0.001*, *corrected*) than those from all unmatched pairs (the red bar), indicating that the group-level CAPs are spatially similar to their matched RSNs. Furthermore, spatial correlation values from all matched pairs are found to be statistically significantly higher (*p<0.001*, *corrected*) than those from positive-only unmatched pairs (the green bar), indicating that matched pairs have the best matches in general while CAPs that are anatomically neighbored do show some overlaps.

**Figure 1.**
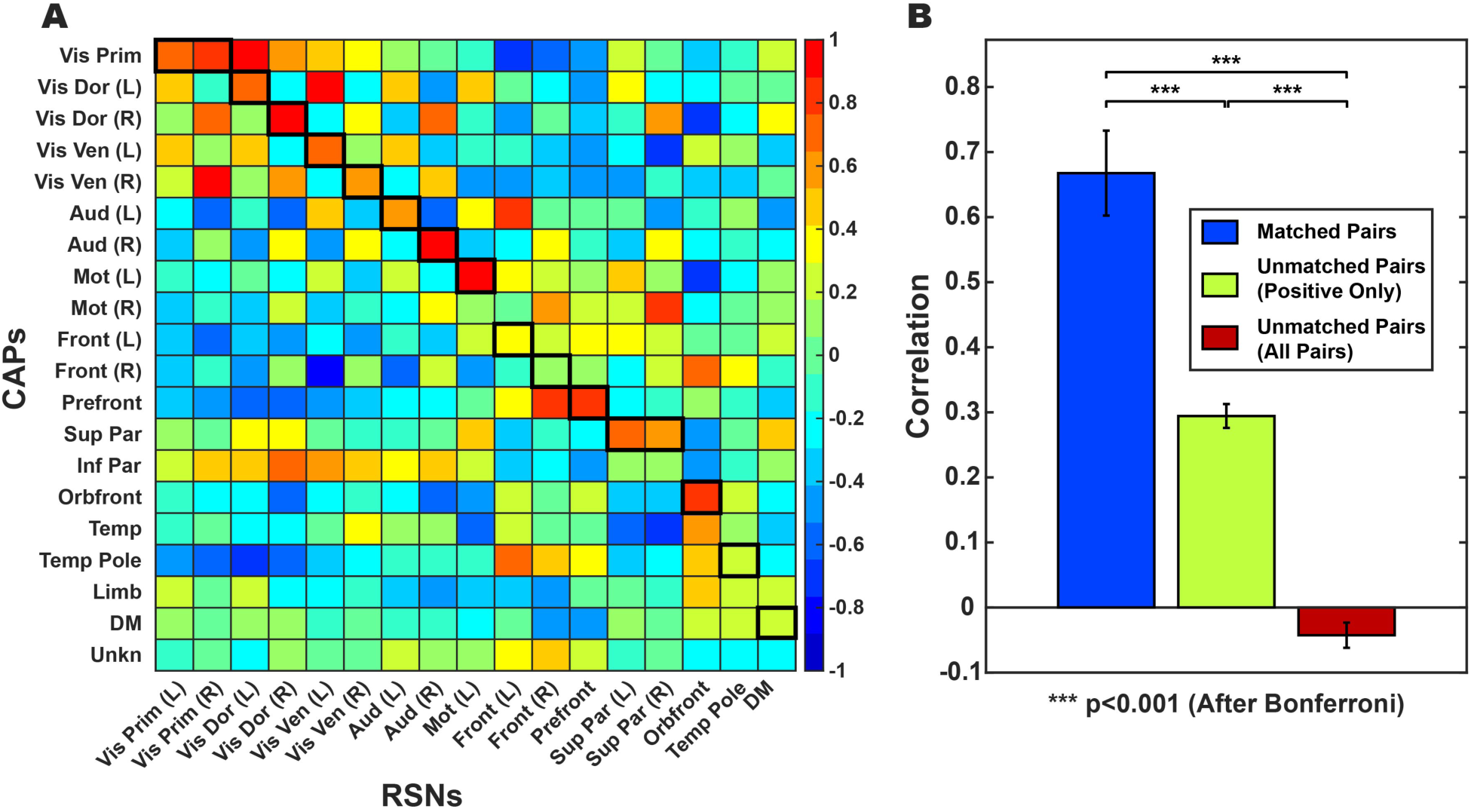
Spatial correlation between CAPs and RSNs. (**A**) The matrix of spatial correlation values (converted to Fisher z-scores) between any pairs of a CAP (out of total 20) and an RSN (out of total 17 after removing duplicated matches). Matched CAP-RSN pairs are boxed in black (see Section 2.5 for the matching protocol). (**B**) Statistical comparisons between spatial correlation values from the matched CAP-RSN pairs and the unmatched pairs after Bonferroni correction. Error bars display standard error of the mean.

The spatial similarities between matched CAPs and RSNs can also be visually observed directly in their spatial maps. Fig. 2 illustrates SM CAPs and their matched RSNs, which anatomically encompass the primary visual network (Vis Prim), dorsal visual stream (Vis Dor), ventral visual stream (Vis Ven), auditory (Aud), and sensorimotor (Mot) networks. Fig. 3 illustrates HO CAPs and their matched RSNs, which anatomically cover the frontal (Front), prefrontal cortex (Prefront), orbital frontal (Orbfront), superior parietal (Sup Par), inferior parietal (Inf Par), temporal pole (Temp Pole), and temporal (Temp) regions. Fig. 4 illustrates Other CAPs, which exhibit spatial patterns similar to DMN and limbic network (Limb), as well as an unknown network (Unkn) that hemispheric-symmetrically runs along the inferior frontal sulcus, cuts through the sensorimotor cortex above the lateral fissure, and then extends the occipito-parieto-temporal junction. Among these CAPs, the corresponding RSN is only identified in the DMN CAP. It is observed via comparing Figs 2, 3, and 4 that SM CAPs display notable similarities to their RSN counterparts while the spatial correspondence of HO and Other CAPs to their matched RSNs is less distinct. This observation is further confirmed by the quantitative correlation values in the confusion matrix (Fig. 1A), where the boxed elements on the left top quadrant (for SM CAPs) show higher values generally than the boxed elements on the right bottom quadrant (for both HO and Other CAPs).

**Figure 2.**
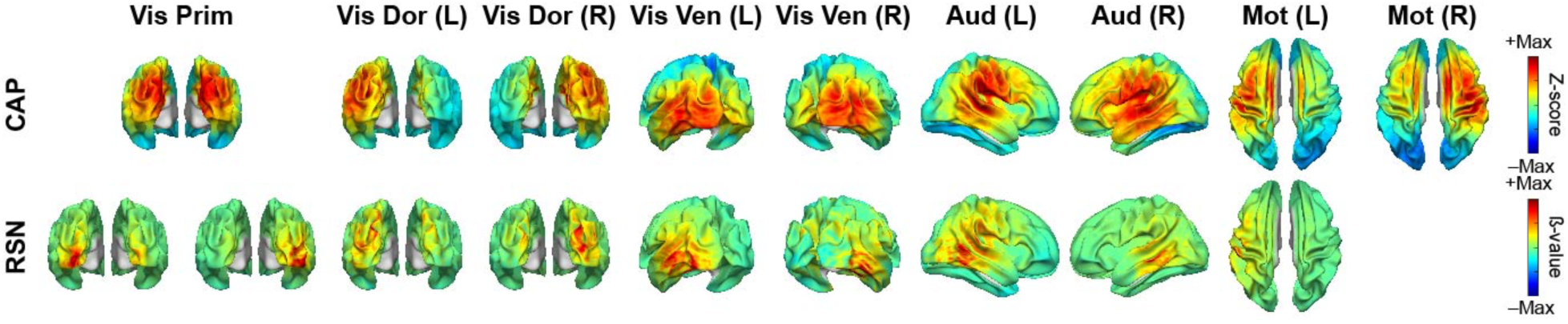
Sensorimotor CAPs and their corresponding RSNs, including primary visual (Vis Prim, matched to two RSNs), visual dorsal stream (Vis Dor), visual ventral stream (Vis Ven), auditory (Aud), motor (Mot, no RSN match for Mot (R)) CAPs. L: left, R: right.

**Figure 3.**
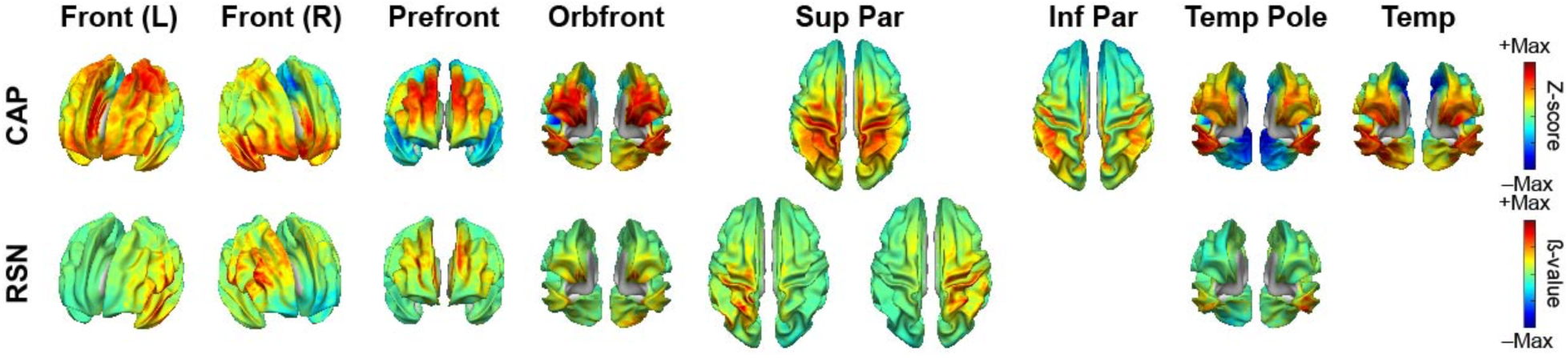
High-order CAPs and their corresponding RSNs, including frontal (Front), prefrontal (Prefront), orbital frontal (Orbfront), superior parietal (Sup Par, matched to two RSNs), inferior parietal (Inf Par, no RSN match), temporal pole (Temp Pole), and temporal (Temp, no RSN match) CAPs.

**Figure 4.**
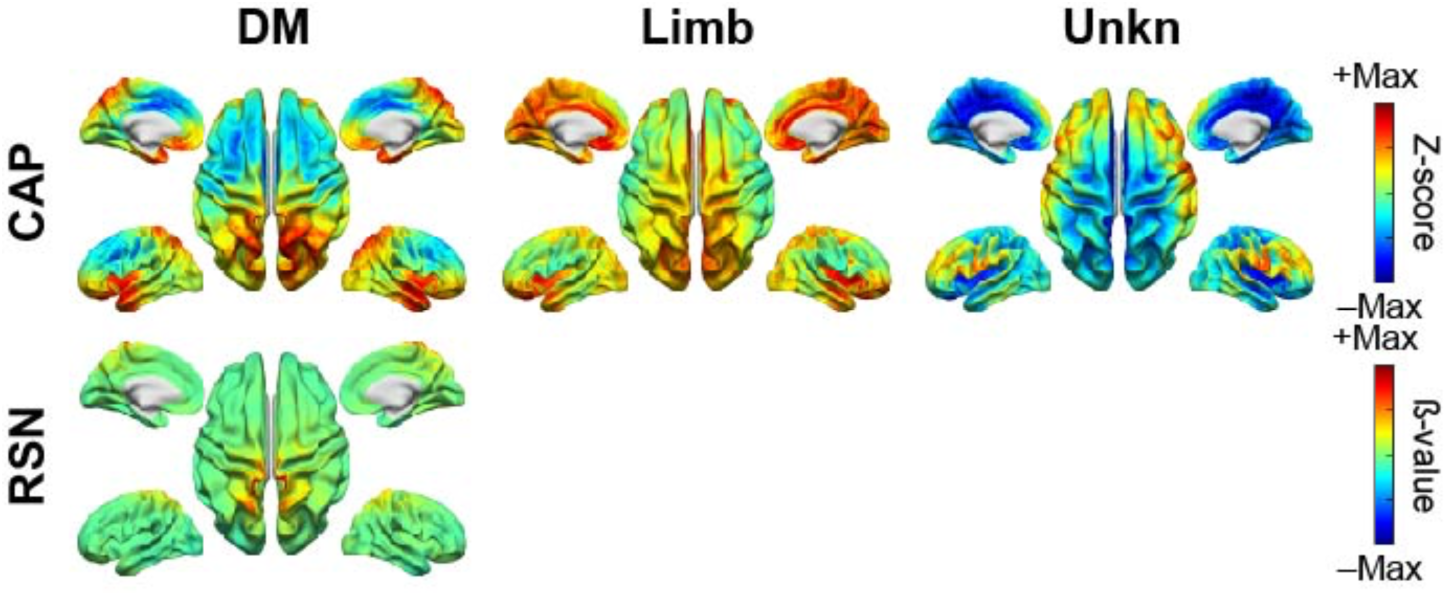
Other CAPs and their corresponding RSNs, including default mode network (DM), limbic (Limb, no RSN match), and unknown (Unkn, no RSN match) CAPs.

The five CAPs that are not matched include one from the SM CAP group, i.e., Mot (R) CAP, and four from both the HO and Other CAP groups, i.e., Inf Par, Temp, Limb, and Unkn CAPs. The hemispheric counterpart for the Mot (R) CAP, i.e., the Mot (L) CAP, is matched to a RSN with a high spatial correlation value (i.e., 0.79).

### 3.2 Hemispheric Symmetries of CAPs

The cortical maps of CAPs exhibit distinct spatial patterns of hemispheric symmetries, which are presented either as pairs of left- and right-dominant unilateral CAPs or individual bilaterally symmetric CAPs. Among the 20 identified CAPs, 10 CAPs are presented as unilateral CAP pairs (i.e., Vis Dor, Vis Ven, Aud, Mot, and Front), while the remaining 10 CAPs are presented as individual bilateral CAPs (Vis Prim, Prefront, Sup Par, Inf Par, Orbfront, Temp, Temp Pole, Limb, DM, Unkn). The quantitative measure of symmetric index supports the strong hemispheric symmetries of CAP cortical maps with the score of 0.89 ± 0.04 for unilateral CAP pairs and the score of 0.91 ± 0.01 for individual bilateral CAPs (Fig. 5B). It is also observed that both HO and Other CAPs appear more often as bilateral spatial patterns than SM CAPs. Only 1 of 9 SM CAPs (i.e., Vis Prim) is bilateral, compared the 6 out of 8 HO CAPs and the 3 of 3 Other CAPs which show bilateral patterns.

**Figure 5.**
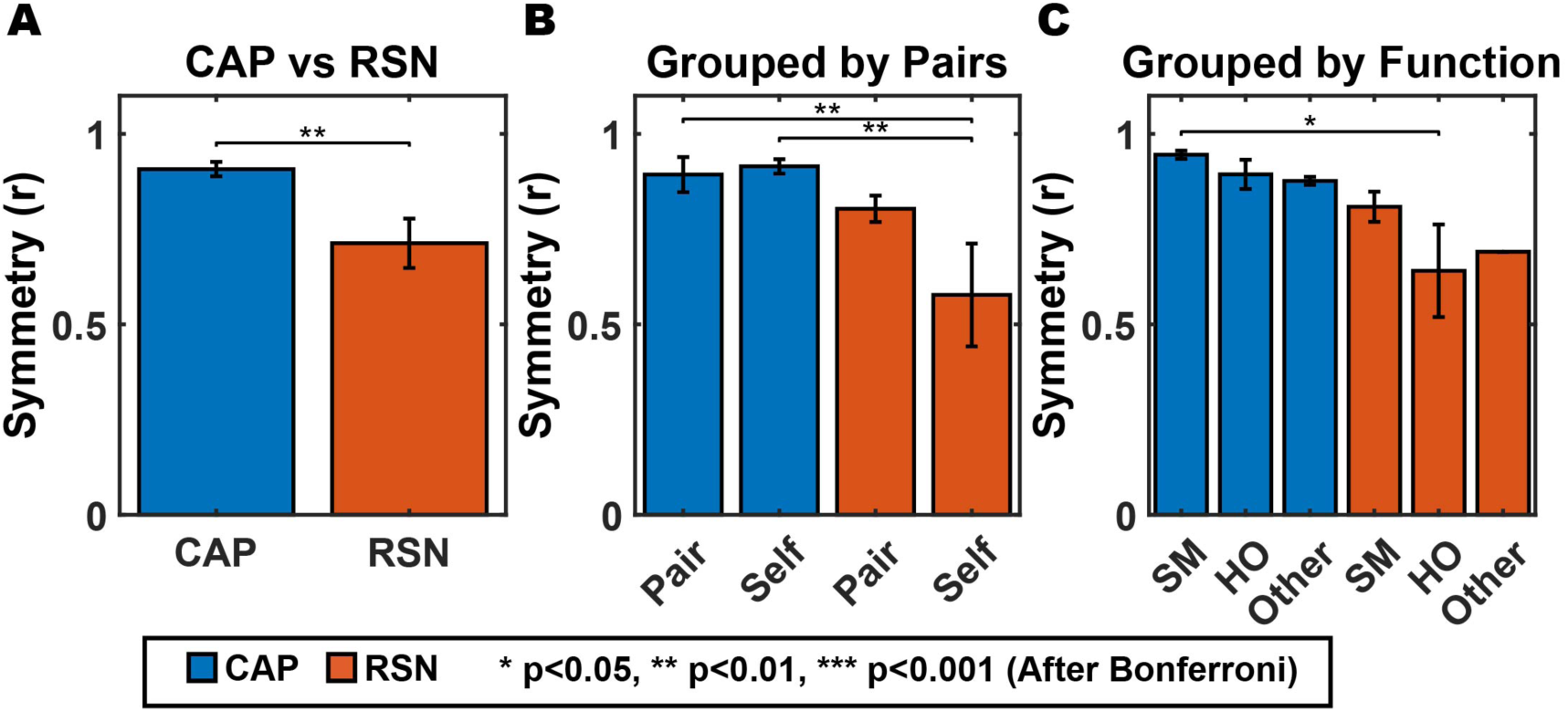
Hemispheric Symmetries of CAPs and ICs. The bar plots display the hemispheric symmetry (r) values of CAPs and ICs grouped by (**A**) overall; (**B**) status of being symmetric pairs (one L CAP/RSN and one R CAP/RSN) or single bilateral CAPs/RSNs (self); and (**C**) by three groups (SM, HO, and Other groups). Error bars represent standard error of the mean.

The cortical maps of RSNs also exhibit hemispheric symmetries while a smaller number of RSNs have bilateral symmetric patterns on both hemispheres (4 out of the 17 ICs). The majority of RSNs (13 of the 17 ICs) appear as pairs of symmetric unilateral CAPs (with the exception of Mot L that was not paired). In comparison to CAPs, the symmetric index values for RSNs significantly decrease overall (*p < 0.01, corrected*) (Fig. 5A), as well as in both unilateral-CAP pairs (*p < 0.01, corrected*) and individual bilateral CAPs (*p < 0.001, corrected*) (Fig. 5B). Meanwhile, the RSNs from the three groups (SM, HO, and Other groups) all show decreased symmetric index values than the three CAP groups, and the decrease in the HO group is the most (Fig. 5C).

### 3.3 Inter-participant consistencies and varibilities of CAP spatial patterns

Fig. 6A shows the boxplots of spatial correlation values between the participant-level CAP cortical maps and their corresponding group-level CAP cortical maps. In general, CAPs display high inter-participant consistencies with their median spatial correlations close to or higher than 0.65 (the majority close to or higher than 0.75) with only one exception (close to 0.55 for the Limb CAP). CAP spatial patterns exhibit certain variabilities among three CAP groups, in which, relatively speaking, the SM CAPs have the highest consistencies, the HO CAPs are in the middle, and the Other CAPs have the lowest consistencies. T-test results indicate that the spatial correlation values from HO CAPs are significantly lower than SM CAPs (*p < 0.001, corrected*) and Other CAPs are significantly lower than both HO CAPs (*p < 0.001, corrected*) and SM CAPs (*p < 0.001, corrected*). This relative pattern can be easily observed in the histograms of spatial correlation values for the CAPs from these three groups (Fig. 6C). These histograms illustrate that the spatial correlation values for SM CAPs have the narrowest distributions, indicating less variabilities while the spatial correlation values for Other CAPs have the relatively widest distributions, indicating more variabilities. The relatively moderate distributions of the spatial correlation values from HO CAPs indicate their higher variabilities than SM CAPs, but lower variabilities than Other CAPs.

**Figure 6.**
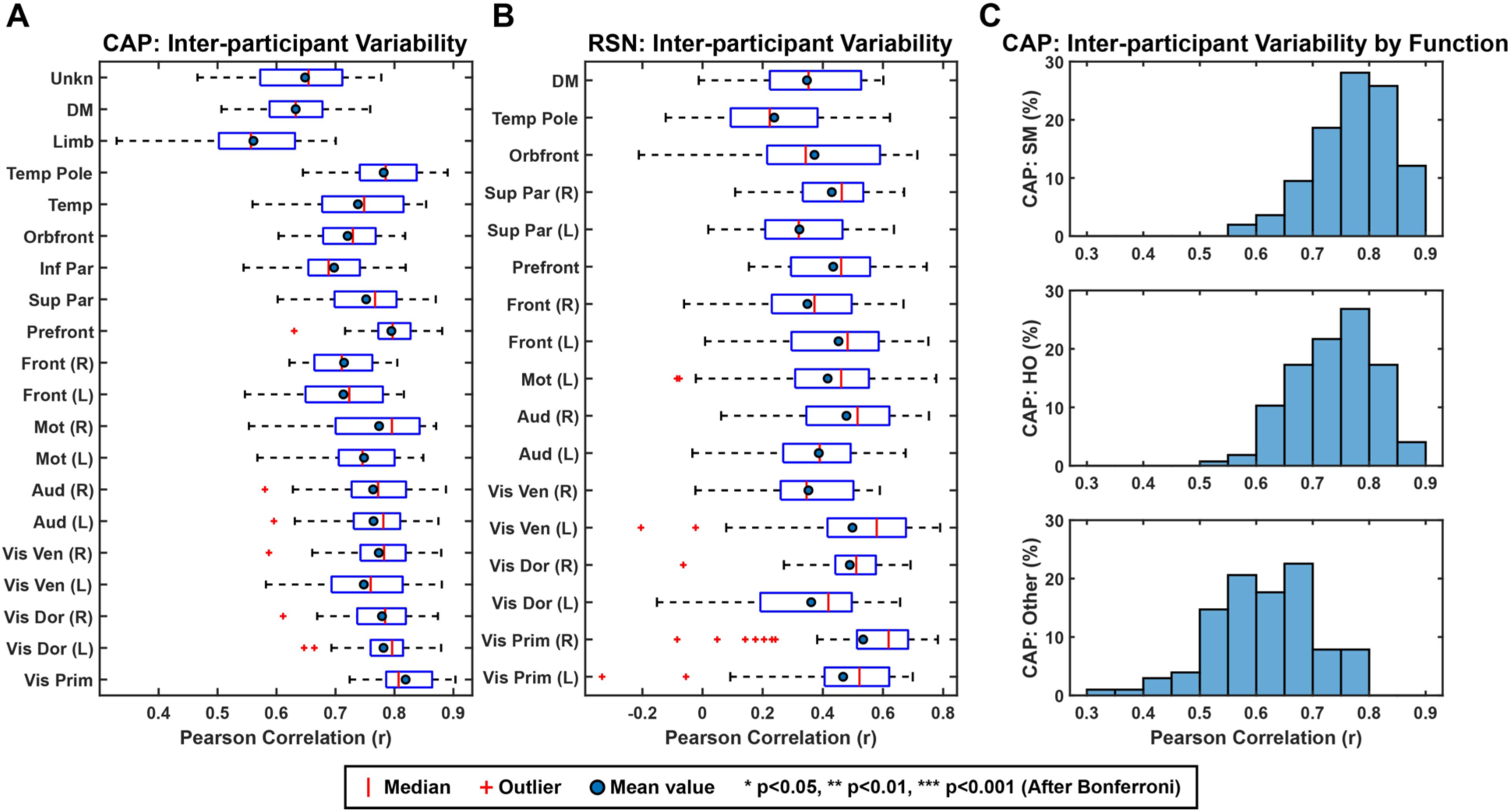
Inter-participant variability of spatial patterns of CAPs and their corresponding RSNs. (**A**) The boxplots of spatial correlations between the participant-level spatial maps and their corresponding group-level spatial maps of 20 CAPs. (**B**) The boxplots of spatial correlations between the participant-level spatial maps and their corresponding group-level spatial maps of 17 ICs. Red bars: median; blue dots: mean. (**C**) Histograms of spatial correlations between the participant-level and group-level spatial maps of 20 CAPs for three group of CAPs (SM, HO, and Other). The y-axis has been converted to percentages to account for the disproportionately lower number of Other CAPs compared to the other two groups.

In contrast, the inter-participant consistencies of RSNs are distinctly lower (Fig. 6B), as all their median spatial correlations close to or less than 0.6 (the majority less than 0.45). To investigate the comparatively low inter-participant consistencies of RSNs, individual-level ß-values obtained during the regression analysis (see Section 2.3) from the top 5% of activated nodes selected from the group-level RSN spatial maps were examined and their mean values from all individuals are visualized using the boxplots (Supplementary Fig. 2B). It is observed that RSNs with higher mean ß-values from all participants usually have greater inter-participant consistencies of spatial patterns (Supplementary Fig. 2A), which is supported by the positive correlation (r = 0.73, *p* < 0.05) between the ß-value magnitudes and the inter-participant correlations from all RSNs and all participants (Supplementary Fig. 2C).

### 3.4 Recurring transient patterns of CAPs

While CAPs show high similarities to time-averaged RSNs in spatial patterns, CAPs temporally are more characterized as recurring transient events. The lifetimes of all 20 CAPs, averaged across all participants, are consistently in the range 66–94 ms (Fig. 7B), demonstrating the sub-second, transient nature. These lifetime data suggest that resting human brains rapidly transition among a set of configurations dominated by various functional systems (i.e., SM, HO, and Other CAPs) rather than dwelling in any one configuration for long time. At the same time, all CAPs recur at the average rates lower than 1 Hz, each of which remain largely consistent across participants (largely varies between 0.5 Hz and 0.9 Hz except the three Other CAPs). It is also noted that all 20 CAPs were present in all individual participants. These observations indicate that these CAP patterns emerge intermittently, and the brain states represented by them are being visited in a repeatable manner over time.

**Figure 7.**
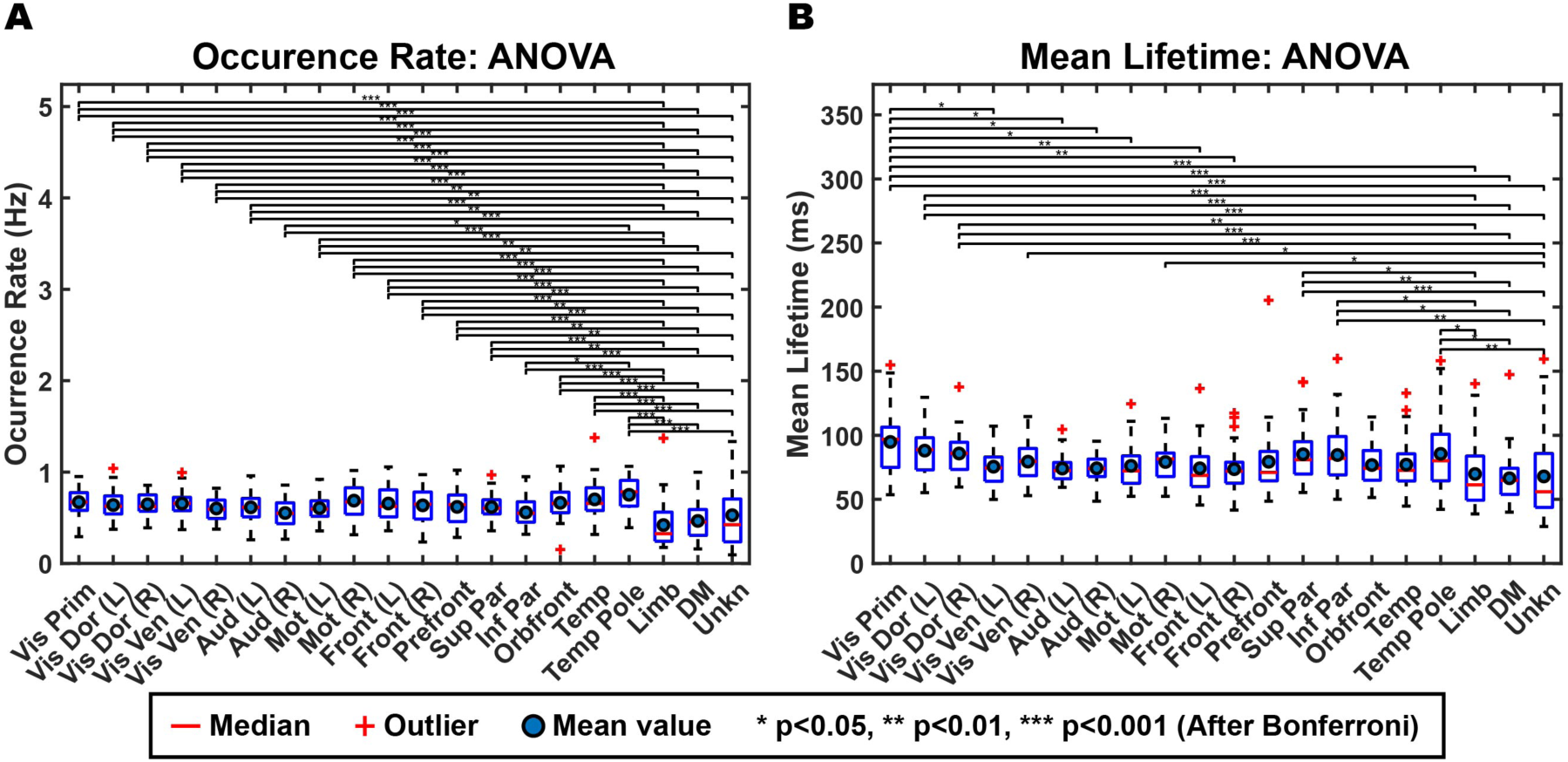
Boxplots of the occurrence rate (**A**) and mean lifetime (**B**) for 20 CAPs. Differences were assessed using ANOVA and then post-hoc t-test on the log-transformed values of occurrence rate and lifetime and statistical significances were corrected using the Bonferroni correction for total 190 pairs.

Among the three groups, the Other CAPs exhibit the largest variabilities in both mean lifetime and occurrence rate among individuals (Fig. 7). One-way ANOVA on the log-transformed occurrence rates and mean lifetimes performed over all possible pairs of 20 CAPs revealed that 47 out of total 49 statistically significant differences (*p < 0.05, corrected*) in occurrence rate and 20 out of total 26 statistically significant differences (*p < 0.05, corrected*) in mean lifetime observed are associated with the three Other CAPs. Moreover, significant differences for occurrence rate at *p < 0.01* (total 47) and mean lifetime measures at *p < 0.001* (total 9) observed are all associated with the three Other CAPs. These significant differences indicate lower occurrence rates of these three CAPs than most of other CAPs, with the Limb CAP exhibiting the lowest occurrence rate (Fig. 7A). The results for lifetime indicate shorter mean lifetimes of these three CAPs than all other CAPs, but the difference only reaches statistically significant levels with three visual CAPs, i.e., the Vis Prim and Vis Dor (L and R) CAPs, which have the longest lifetimes (Fig. 7B). These three CAPs further exhibit increased inter-participant variability compared to all other CAPs (e.g., increased heights of boxes in Figs. 7A-B indicating the difference between 75^th^ percentile and 25^th^ percentile).

### 3.5 Reproducibility of CAPs due to k

Beyond the reproducibility of CAP’s spatial (Fig. 6A) and temporal patterns (Fig. 7) from individuals, the reproducibility of CAPs due to pre-selected parameter for the k-means algorithm, i.e., the number of clusters (k), was investigated. Fig. 8 illustrates the group-level cortical maps of CAPs with the k value increases from 10 to 20. In general, identified CAPs at different k values remain spatially consistent, demonstrating their reliable detection even when the k value changes in a relatively wide range (i.e., doubled). When the k value increases, additional clusters emerge as subdivisions from the existing CAPs at low k values. A representative example is the Mot CAP, which shows a bilateral pattern over two hemispheres at k = 10 and starts to split into one left-hemisphere-dominant CAP and one right-hemisphere-dominant CAP from k = 12. Other new CAPs, introduced by increasing k, tend to reflect finer sub-systems within the same functional system. For example, the Vis Prim CAP starts to appear when k = 15 that might be split from both left and right Vis Dor CAPs as their spatial patterns shift slightly lateral at k = 15 from k = 12. Both left and right Vis Ven CAPs emerge at k = 18 that might be split from both left and right Vis Dor CAPs as their spatial patterns shift slightly along the dorsal direction and become close to the parietal cortices at k = 18 in comparison to k = 15. Lastly, it is noted that some new CAPs further emerge at the highest k value (i.e., k = 20), which might suggest that further investigations into even higher k values (> 20) might be still valuable. For example, the Temp CAP only appears at k = 20 probably due to splits from both the prefront CAP and the Temp Pole CAP due to its spatial similarities to them. It is also interesting to note that among three Other CAPs, both the Limb CAP and Unkn CAP are consistently present from k = 10 to 20 while the DMN CAP only shows up when k = 20. These examples not only indicate the persistence of these patterns across different k values but also suggest that CAPs are not merely artifacts of a specific clustering choice but instead reflect functional systems that can be resolved at multiple levels of granularity.

**Figure 8.**
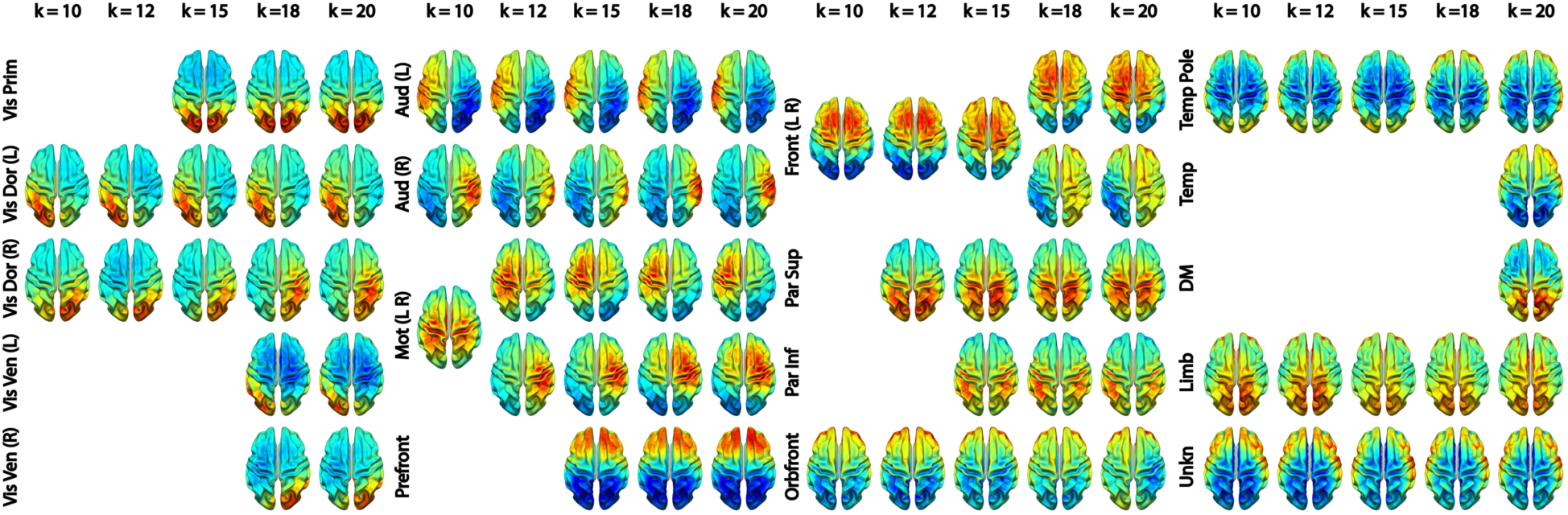
Spatial Patterns of CAPs for k = 10, 12, 15, 18, and 20.

## 4. Discussion

In the present study, we identified a set of reproducible recurring and transient co-activation patterns (CAPs) in EEG that resemble canonical time-averaged RSNs. Beyond spatial matches, CAPs exhibit RSN-like characteristics, including hemispheric symmetry, separatable granularity in functional systems, intersubject variability gradient across low-level and high-level functional systems, and the existence of both DMN and TPNs. These findings suggest that time-averaged RSNs, constrained due to the use of time-domain measures (e.g., correlation and independence), might be expressions of brief, recurring, spatially large-scale neuronal events. On the contrary, CAP analysis utilizing spatial-domain measures is able to resolve such spatially large-scale neuronal events at much fine temporal scales (<100 ms), which potentially offers a new window on investigating fast dynamics of such large-scale neuronal events.

### 4.1 Spatial Patterns of CAPs

Our findings provide the evidence that cortical CAPs reconstructed from EEG data exhibit strong spatial similarities to time-averaged RSNs that have been widely reported in fMRI literatures (Beckmann et al., 2005; Damoiseaux et al., 2006). Nine SM CAPs recover distributed cortical co-activation patterns within the somatosensory, visual, auditory, and motor systems. For the visual system, the existence of multiple CAPs indicates that the primary, ventral, and dorsal visual cortices are spatially separatable in the functional networks defined by CAPs. Two unilateral motor CAPs reveal symmetric but opposite hemisphere-dominant distributed activations each involving both the primary motor cortex and supplementary motor areas (Fig. 2). The DMN CAP reveals the full membership of cortical nodes of DMN established in fMRI within the medial prefrontal cortex, posterior cingulate cortex, temporo-parietal junction, and temporal lobes on both hemispheres. Several HO CAPs (left and right Front, Prefront, and Orbfront CAPs) cover the areas for executive control networks identified in fMRI (Beckmann et al., 2005). Beyond the fMRI literature, the spatial patterns of these CAPs are also similar to time-averaged RSNs found in EEG data (Q. Liu et al., 2017; Yuan et al., 2016) and MEG data (Brookes et al., 2011; De Pasquale et al., 2010; Hipp et al., 2012), which is not surprising as the spatial correspondences between EEG RSNs and fMRI RSNs have been reported (Yuan et al., 2016). Such similarities are also investigated in the present study and our results support the strong correspondences between CAPs and RSNs derived from the same EEG dataset (Figs. 2-4), which is further supported by high spatial correlations (of statistical significance) in all matched CAP-RSN pairs (Fig. 1B). The similarities between CAPs and RSNs are not only observed among individual matched pairs, but also collectively among the entire identified sets of CAPs. In the present study, of the 20 CAPs identified, 15 CAPs show high similarities to RSNs reconstructed from the same dataset. Compared to the fMRI literature, the only frequently reported network patterns missing from the set of CAPs are the frontoparietal RSNs (Damoiseaux et al., 2006), which are however also often missed in EEG/MEG RSNs (Q. Liu et al., 2017). Considering the detections of both frontal and parietal CAPs (Fig. 3), a potential hypothesis is that these networks might prevail in different frequency ranges other than the alpha band investigated in the present study as large-scale synchronizations are frequency-specific (Hipp et al., 2012; Vidaurre et al., 2018).

The present study also provides evidence that suggests the correspondence between CAPs and RSNs beyond their matching spatial patterns. Firstly, CAP spatial patterns display high hemispheric symmetry (Fig. 5) and strong hemispheric symmetries are the hallmark of RSNs (Smith et al., 2009) and other network patterns from fMRI such as resting-state functional connectivity (Calhoun et al., 2014), CAPs (Gutierrez-Barragan et al., 2019; X. Liu et al., 2013), and large-scale propagation waves (Matsui et al., 2016; Mitra & Raichle, 2016). Such hemispheric symmetries are also observed in electrophysiological (Takeda et al., 2021; Yuan et al., 2016) and optical studies (Khan et al., 2021, 2022, 2024) regarding RSNs, CAPs, and other large-scale resting-state activations and/or phenomena, as well as in EEG RSNs from the present study (Fig. 5). Secondly, the functional systems represented by CAPs could be separated into multiple levels of granularity depending on chosen clustering parameters. For example, varying the k value from 10 to 20 illustrates the process of generating CAP representations for more visual sub-systems within the visual cortex (Fig. 8). The separation of the motor system from a bilateral CAP at k = 10 to two hemispheric-dominant unilateral CAPs at k = 12 is another example (Fig. 8). These observations are consistent with hierarchical anatomic organizations of human cortical structures (Thomas Yeo et al., 2011). Their separatable representations in functional data have been reported in fMRI where more RSNs appear to represent sub-systems when the complexity of model increases, e.g., the number of ICs (Smith et al., 2009). Lastly, increased inter-participant variability is observed from SM CAPs to HO CAPs (Fig. 6A & 6C), which is consistent with intersubject variability gradients (Mueller et al., 2013; Seitzman et al., 2019) across lower-order perceptual networks (lower inter-participant variability) and higher-order functional networks (higher inter-participant variability). These observations suggest that the spatial patterns of CAPs are modulated in the same direction as RSNs on three important properties of functional neural systems, i.e., homologue, sub-systems, and individual variability, therefore indicating possible common distributed neuronal substrates behind both CAPs and RSNs.

Several large-scale patterns of transient nature (at similar timescales to CAPs) have been identified from EEG/MEG, i.e., microstates (Britz et al., 2010; Custo et al., 2017; Pascual-Marqui et al., 2014), HMM (Baker et al., 2014), and CAPs based on a different distance measure, i.e., L1-norm (Ding et al., 2022; Shou et al., 2022). Many of them have been interpreted in the context of RSNs based on their spatial similarities to RSNs. For example, convolving time courses of microstate with hemodynamic response functions (HRFs) have yielded RSN-like spatial patterns (Britz et al., 2010). Similarly, HMM-derived brain states from MEG have been found spatially similar to canonical RSNs (Baker et al., 2014). L1-norm distance-based CAPs reveal several RSN-like configurations (e.g., visual, sensorimotor, and DMN networks) when IC time courses are used for clustering (Ding et al., 2022). On the other hand, differences are also observed among these transient patterns. Traditional microstate analyses often only resolve four dominant patterns (Michel & Koenig, 2018) each with a dipolar topography that is usually generated by focal regional activations. Moreover, cortical activation patterns identified with microstate and HMM analyses have been indicated with significantly different spatial patterns (Coquelet et al., 2022). L1-norm distance-based CAPs reveal unique global coactivation patterns that have not been reported in any other transient large-scale phenomena (Ding et al., 2022). At last, L1-norm distance-based clustering algorithm on time courses of anatomical cortical parcels (Shou et al., 2022) seems to reconstruct a set of CAPs of almost completely different spatial patterns (revealing more between-CAP differences in terms of global magnitudes) as compared with the CAPs obtained using the correlation distance in the present study (revealing more between-CAP differences in terms of spatial structures). Such similarities and differences among various transient brain-wide activation patterns might be explained due to the existence of multiple large-scale neuronal activation mechanisms and varying sensitivities of different models/algorithms to different activation mechanisms. The significant differences between CAPs obtained using different distance measures, i.e., L1-norm vs correlation, present the strong evidence in supporting the hypothesis as the sensitivity to detect different large-scale activations could be switched with the only change of a key parameter but using the same approach (i.e., clustering).

### 4.2 Temporal Characteristics of CAPs

EEG CAPs obtained in the present study exhibit two distinct temporal characteristics: transience and recurrence. Brain states represented in CAPs manifest themselves as recurring transient episodes of large-scale co-activations between 60 and 100 ms across all participants. All brain states (i.e., 20 CAPs) are visited by every participant, with stable occurrences and lifetimes across all participants and CAPs.

Prior work in EEG/MEG supports the presence of transient brain states lasting on the order of tens to hundreds of milliseconds. Microstate analyses have repeatedly shown that EEG topographies remain quasi-stable for durations below 200 ms (Pascual-Marqui et al., 2014; Custo et al., 2017). HMM analyses on MEG data identify short-lived states on the order of 50–200 ms (Baker et al., 2014; Vidaurre et al., 2018). L1-norm based CAPs exhibit their lifetimes between 25 to 40 ms (Ding et al., 2022; Shou et al., 2022). While variations exist in various large-scale transient neuronal activations, their timescales are consistent with neuronal events revealed in evoked brain activations (tens to hundreds of milliseconds) (Light et al., 2010). Therefore, these lifetime values are potentially indicative of the actual lifetimes of underlying neuronal events, adding new understanding to large-scale networked activations beyond time-averaged RSNs. It is also noted that the selection of approach parameters might impact this temporal metric as well. Our results indicate that, as the k value increases from 10 to 20, the mean lifetimes of CAPs decrease from 102–129 ms to 66–94 ms. Furthermore, all large-scale transient neuronal activations discussed above (i.e., microstates, HMM, and CAPs with various distance measures) exhibit recurring behaviors.

Our results further suggest the CAPs from the Other group show significantly shorter lifetimes and lower occurrence rates than many CAPs from both the SM and HO groups. Such significant differences are not observed among any pair of CAPs from within either the SM or HO group or from across these two groups. Shortened lifetimes and lowered occurrences of the Other CAPs might lead to their insufficient and unequal representations within the total 10-minute recordings in individual participants that can explain their significantly higher inter-participant variabilities of spatial patterns (Fig. 6A). Furthermore, the inter-participant variabilities of these two temporal measures in the Other CAPs seem also relatively higher than many CAPs from both the SM and HO groups (Fig. 7), which supports the higher inter-participant variabilities of these CAPs. As the three Other CAPs indicate either highly networked activations or activations outside sensory and motor cortices, these observations are consistent with intersubject variability gradients (Mueller et al., 2013; Seitzman et al., 2019) across lower-order perceptual networks and higher-order functional networks as discussed above.

### 4.3 Relationship between CAPs and RSNs

Our results and observations seem to support the hypothesis that the canonical time-averaged RSNs may stem from temporal-aggregations of repeated transient episodes of co-activations (i.e., CAPs). Firstly, the majority of CAPs and RSNs display significantly similar spatial maps (Figs. 1-4). The CAPs that could not be spatially matched to RSNs (e.g., the Other CAPs in Figs. 6 and 7) typically exhibit shorter lifetimes, lower occurrences, and greater inter-participant variability, all of which reduce the formation of spatially and temporally stable patterns over entire recording windows in individual participants and across all participants, thereby potentially limiting corresponding reliable component extraction by ICA. Secondly, while individual CAPs seem transient, most of them consistently recur throughout entire recording windows and all participants. Their repeated activations provide temporally sustained components that might lead to their presences as time-averaged RSNs. This is further supported by the observations that the CAPs with high occurrences (e.g., visual, auditory, temporal) are better spatially matched to time-averaged RSNs, whereas the CAPs with low occurrences (e.g., the Limb, Unkn CAPs) are less spatially matched or not matched (Fig. 1A). Lastly, both CAPs and RSNs largely show hemisphere-symmetric patterns (Figs. 2-5) that offer additional evidence about their potential same underlying neural generators. The hypothesis between CAPs and RSNs is supported by several reported fMRI findings. Liu and Duyn (2013) shows that averaging the top 1–15% of timeframes based on fMRI signal amplitudes from pre-selected seeds produces spatial maps highly similar to RSNs derived from a full correlation analysis. Related work further emphasizes that RSN structures could be recovered from high-activation time points alone, without requiring stationary assumptions (Gutierrez-Barragan et al., 2019; Tagliazucchi et al., 2012). While fMRI and EEG are in extremely different timescales, these literature support the hypothesis that long-timescale neuroimaging patterns may emerge from the recurrence of short-lived, structured brain events. Taken together, CAPs seem to offer a time-resolved perspective on the same large-scale functional architecture behind RSNs.

### 4.4 Reproducibility

CAPs are consistently reproduced as stable patterns across a range of the number of clusters (i.e., k=10-20). It also meets the expectation that increasing the number of clusters results in revealing more hierarchical structures in neural systems (e.g., the visual system in Fig. 2), which is the phenomenon similarly observed in fMRI data as discussed above. To further assess the reproducibility of CAPs, four additional iterations of clustering with k=20 were run using randomly generated seeds as the k-means clustering is an iterative algorithm. Resulting CAPs from different iterations could be well matched to the CAPs in the primary analysis (see examples in Supplementary Fig. 3). It is noted that minor variations are observed in the Limb and Front (L, R) CAPs across iterations. The Limb CAP alternates between localizing to the cingulate cortex and extending to broader areas on the middle wall across iterations. The Front CAPs vary slightly in their degree of symmetry between the left and right hemispheres across iterations. Sensorimotor and DMN CAPs, by contrast, show no noticeable variations across iterations (data not shown), consistent with their low inter-participant variability and robust group-level presence.

The reproducibility is also attested in relatively low to moderate inter-participant variabilities in the spatial patterns (Fig. 6A) and temporal measures (Fig. 7) of all CAPs. It is noted that, in comparison to SM and HO CAPs, the Other CAPs are less consistently detected across participants. However, despite occurring less frequently, lasting much shorter, appearing less consistently across participants, and in general accounting for a much smaller amount of the EEG data, two of the Other CAPs (e.g., Limb and Unkn) exhibited highly reproducible spatial structures across clustering iterations, appearing as early on as k = 10 (Fig. 8).

### 4.5 Methodological Considerations and Limitations

The key methodological difference between the approaches in reconstructing RSNs and CAPs lies in the different domains where similarity measures are calculated. Correlation and independence are time-domain measures, which utilize time samples to measure similarities among different spatial locations (voxels in fMRI and nodes in EEG cortical tomography), and therefore must be calculated over a time window (typically in tens of seconds to tens of minutes dependent on the factors such as nature of contrast signal, sampling rate, research question being asked, and etc.). On the contrary, the distance measures used to obtain CAPs are defined in the spatial domain that calculate similarities among data from individual timeframes. It turns out that CAPs can capture transient events with lifetimes (<100 ms) several orders shorter than the typical time window sizes used for calculating time-domain measures. While the present study and our previous studies (Ding et al., 2022; Shou et al., 2022) demonstrate the CAPs in EEG, similar transient events in fMRI using the concept of CAPs have also been successfully demonstrated (X. Liu et al., 2013). Furthermore, the present study together with our previous studies (Shou et al., 2022) further reveal that the use of different distance measures in the same clustering algorithms can lead to detection of different large-scale transient neuronal patterns, which suggest that there are potentially multiple neuronal processes ongoing at the same time. In contrast, the uses of different time-domain measures (e.g., correlation vs independence) typically lead to similar spatial constructs of RSNs (Petersen et al., 2000). Multiple neuronal processes also address the gap of missing spatial similarities between transient CAPs (Shou et al., 2022) and time-averaged RSNs both from EEG data (Yuan et al., 2016) while high spatial similarities exist between transient fMRI CAPs and fMRI RSNs (Liu and Duyn, 2013). Our present results indicate such spatial similarities also exist between EEG CAPs and EEG RSNs, have been overlooked mainly due to a methodological parameter, not due to different nature of contrast signals (EEG vs fMRI).

One of the key assumptions in the k-means algorithm is that each individual timeframe belongs exclusively to one cluster (i.e., one CAP). This temporal exclusivity restricts the ability to model temporally overlapping or concurrent neural processes. Moreover, the averaged spatial patterns of CAPs (both at individual and group levels) should only reflect the dominant neuronal co-activation patterns from all timeframes assigned to the same CAP and differences of overlapping components between individual occurrences of that same CAP would have most likely been filtered out during averaging. Furthermore, it is reasonable to assume that some timeframes within entire resting EEG recordings might only contain noise-like patterns especially considering the transient nature of neuronal events being detected. Their “hard” assignments to individual clusters might add biases on estimating values for temporal measures (e.g., lifetime). Future studies need to consider how to add a cluster for “noise-like” timeframes to address such biases. However, despite temporal constraints of exclusivity, CAPs are not restricted spatially as the same set of brain regions can be involved in multiple CAPs. Actually, several fMRI studies have argued that fMRI CAPs could be considered as the combination or partial decomposition of time-averaged RSNs (Gutierrez-Barragan et al., 2019; Karahanoğlu & Van De Ville, 2015; Mawla et al., 2023). Our EEG CAPs in the present study exhibit similar phenomena of combinations and partial decompositions, for example in the visual systems (Fig. 2) and in the spatially hierarchical subsets of CAPs revealed by varying k values (Fig. 8). Similar methodological considerations on constraining either spatial or temporal domain also exist in methods for estimating time-averaged RSNs, where enforcing independences to either spatial or temporal dimension of data have led to spatial ICA and temporal ICA (Calhoun et al., 2001). When underlying neuronal activations are not strictly separable in both space and time, these methods yield suboptimal separation. Therefore, it is important to understand that results from different approaches might reflect different aspects of underlying neuronal activations and need to be interpreted accordingly.

## Data and Code Availability

Data used in this study are not publicly available due to data sharing restrictions from the IRB but are available from the corresponding author through a data use agreement upon reasonable request.

## Author Contributions

KN: Investigation, Formal Analysis, Methodology, Writing - original draft. HY: Conceptualization, Study Design, Writing - original draft, Funding Acquisition. LD: Conceptualization, Study Design, Methodology, Formal Analysis, Funding Acquisition, Supervision, Writing - original draft.

## Acknowledgments

We thank Dr. Guofa Shou, Dr. Chuang Li, Mr. Junwei Ma, Dr. Yafen Chen, and Dr. Fan Zhang for data collection and Dr. Guofa Shou for developing a draft version of codes.

