## Supplementary Materials for "Recurring Transient Brain-Wide Co-Activation Patterns from EEG Spatially Resembling Time-Averaged Resting-State Networks"

### Supplementary Material

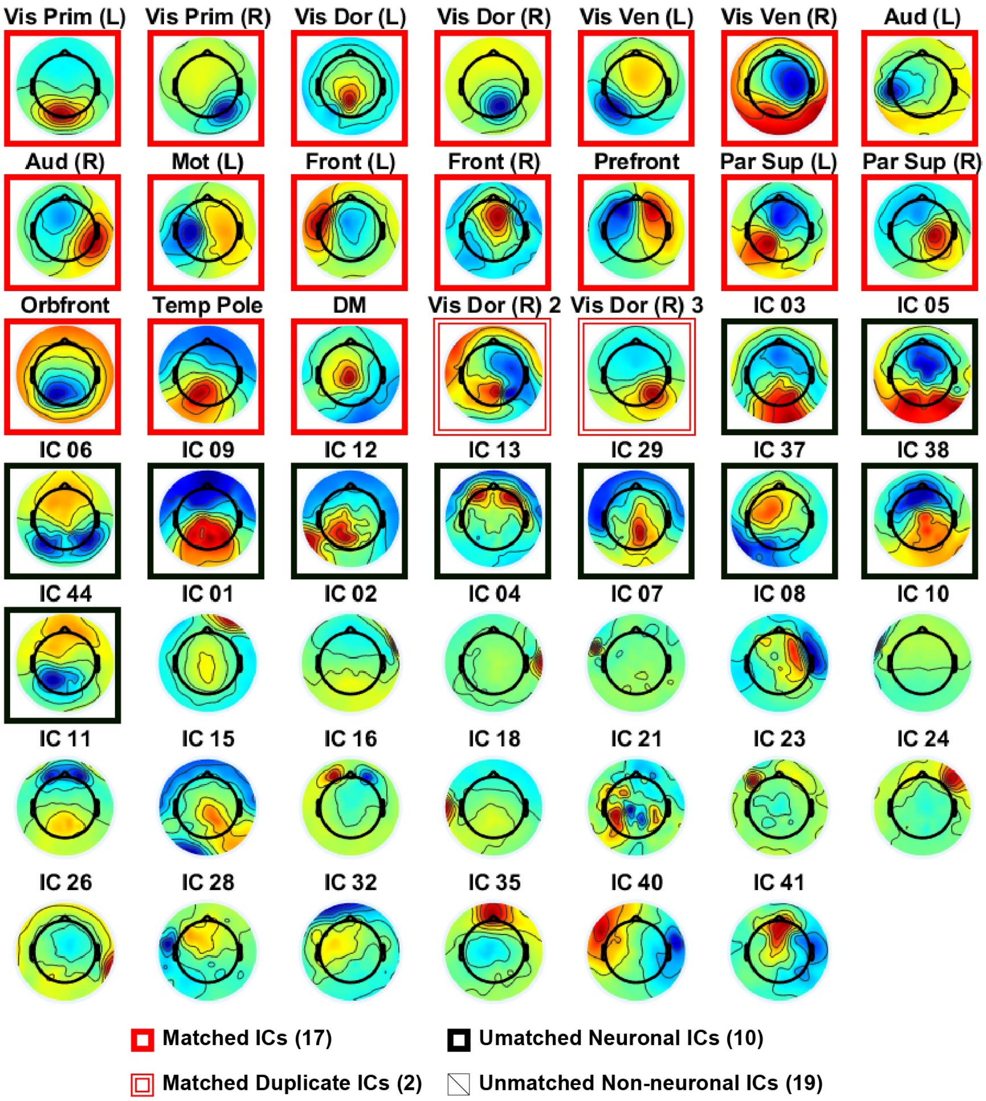


**Supplementary Figure 1.** Scalp Topographies of 48 ICs. All 29 neuronal ICs, identified based on their scalp topographies, temporal and spectral characteristics, are outlined. Red solid outlines: 17 neuronal ICs as matched RSNs to CAPs; red hollow outlines: 2 duplicates of neuronal ICs as matched RSNs; black solid outlines: 10 neuronal ICs as unmatched RSNs.

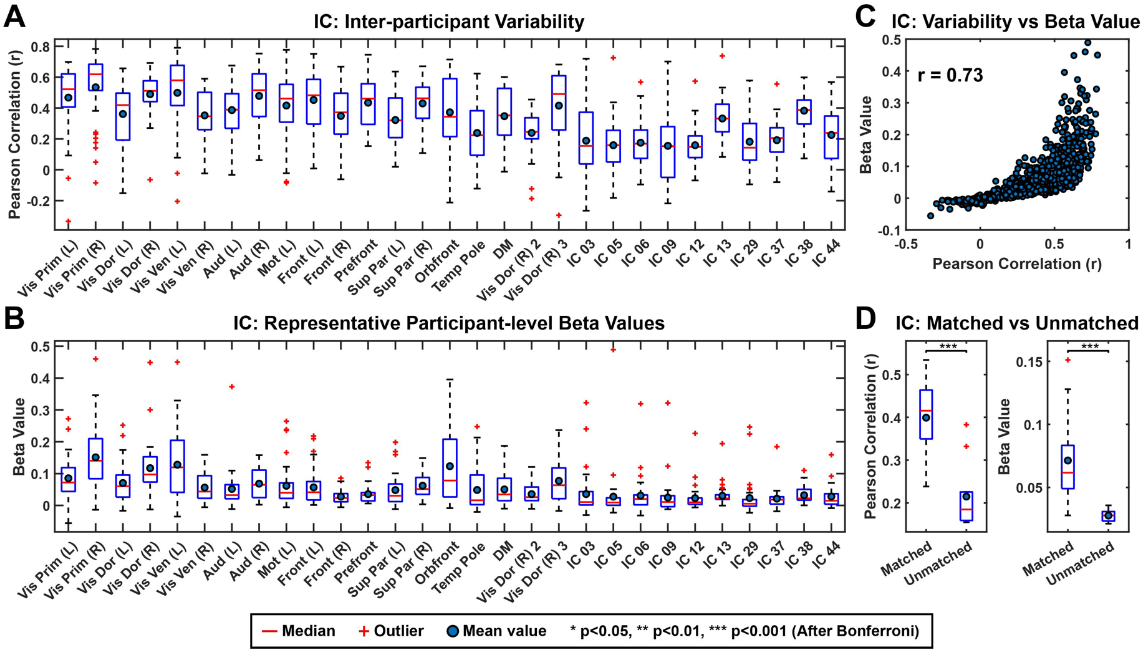


**Supplementary Figure 2.** Correspondence between the inter-participant variability of RSNs (A) and individual-level regression beta coefficients calculated against group-level ICs (B). Sample means are shown by blue bots. (**A**) Box plots of inter-participant variability of all 29 time-averaged RSNs (from 29 neuronal ICs), measured as the spatial correlation of all participant-level cortical tomography of a RSN with its corresponding group-level cortical tomography. The 19 matched RSNs (including two duplicates) to CAPs are named and the remaining 10 unmatched RSNs are labelled with their original IC indices from the output of ICA analysis. (**B**) Box plots of representative individual-participant beta values for all RSNs, calculated as the averages from participant-level cortical tomography over the cortical elements of the top 5% magnitudes determined from the group-level cortical tomography of corresponding RSNs. (**C**) Correlation between the representative beta values of all RSNs from all participants and their corresponding inter-participant variability (measured as Pearson correlation in **A** of this figure). Results show that the ICs with greater participant-level beta values typically exhibit greater inter-participant similarities. (**D**) Box plots of the mean representative beta values (calculated over all participants) and inter-participant variability (measured as Pearson correlation) for the 19 matched and 10 unmatched neuronal RSNs (i.e., ICs). Matched RSNs exhibit statistically significant higher values in both measures than unmatched RSNs, which indicate higher contributions of matched group-level ICs toward variances at participant-level data and greater inter-participant similarities in spatial patterns, respectively.

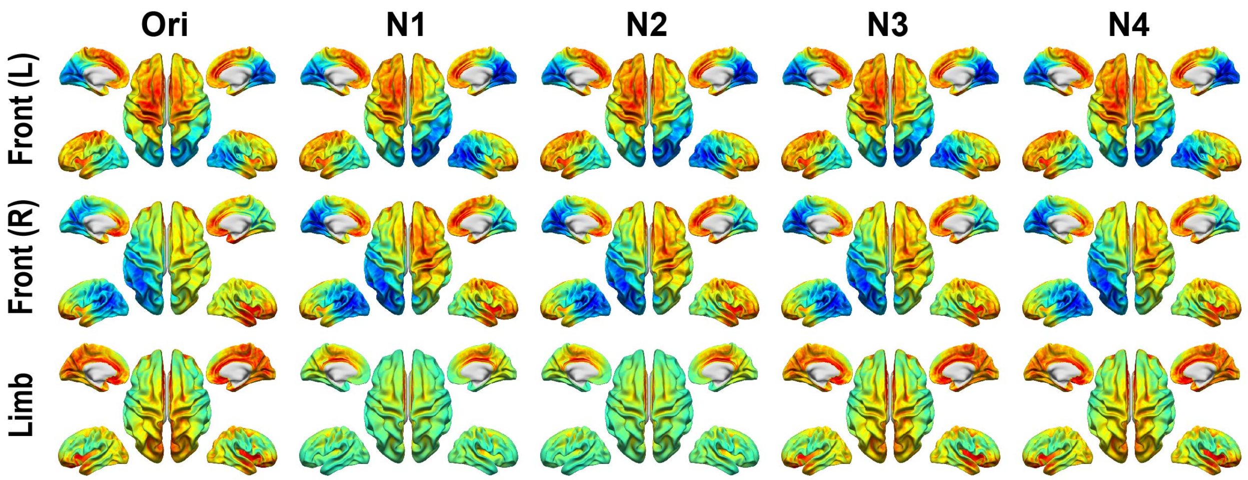


**Supplementary Figure 3.** Alternative clustering realizations based on random seeds for initiating the k-mean algorithm for the Front and Limb CAPs as examples. Original clustering results show a less symmetric pair of Front (L & R) CAPs than clustering results from other four runs. The original clustering results in the Front (L) CAP seem capturing more activations along the superior frontal gyrus on the right hemisphere than the clustering results from other four runs. At the same time, the same activations seem appear more on the Front (R) CAP from other four runs than the original clustering results. The Limb CAP from two runs shows localizing along the middle wall, superior temporal gyrus, and precuneus while the Limb CAP from other three runs localizes within the entire cingulate cortex. Examples for the CAPs from the SM groups are not included here since they exhibit much similar spatial patterns across different realizations. "Ori" refers to the original clustering results presented in the Results section, "N1–N4" refers to the alternative clustering results from other four clustering realizations.
